# Geographic variation in adult and embryonic desiccation tolerance in a terrestrial-breeding frog

**DOI:** 10.1101/314351

**Authors:** T.S. Rudin-Bitterli, J.P. Evans, N.J. Mitchell

**Affiliations:** School of Biological Sciences, The University of Western Australia, Crawley, Western Australia 6009, Australia; Centre for Evolutionary Biology, The University of Western Australia, Crawley, Western Australia 6009, Australia

**Keywords:** amphibians, desiccation tolerance, intra-specific variation, climate change, adaptation, phenotypic plasticity, environmental sensitivity

## Abstract

Intra-specific variation in the ability of individuals to tolerate environmental perturbations is often neglected when considering the impacts of climate change. Yet this information is potentially crucial for mitigating any deleterious effects of climate change on threatened species. Here we assessed patterns of intra-specific variation in desiccation tolerance in the frog *Pseudophryne guentheri*, a terrestrial-breeding species experiencing a drying climate. Adult frogs were collected from six populations across a rainfall gradient and their dehydration and rehydration rates were assessed. We also compared desiccation tolerance of embryos and hatchlings originating from within-population parental crosses from four of the six populations, where selection on desiccation tolerance should be especially strong given that embryos cannot move to escape unfavourable microclimates. Embryos were reared on soil at three soil-water potentials, ranging from wet to dry (ψ = −10, −100 & −400 kPa), and their desiccation tolerance was assessed across a range of traits including survival, time to hatch after inundation, wet mass at hatching, hatchling malformations and swimming performance. We found significant and strong patterns of intra-specific variation in almost all traits, both in adults and first generation offspring. Adult frogs exhibited clinal variation in their water balance responses, with populations from drier sites both dehydrating and rehydrating more slowly compared to frogs from more mesic sites. Similarly, desiccation tolerance of embryos and hatchlings was significantly greater in populations from xeric sites. Taken together, our findings suggest that populations within this species will respond differently to the regional reduction in rainfall predicted by climate change models. We emphasise the importance of considering geographic variation in phenotypic plasticity when predicting how species will respond to climate change.

## INTRODUCTION

Understanding how organisms will respond to rapid environmental change is a major challenge for conservation and evolutionary biologists (Hoffmann and Sgro 2011). Most commonly, projections of climate change impacts on communities and species have been based on studies of the effects of environmental change on single populations (Moran et al. 2016). However, species are not uniform entities (Bolnick et al. 2003, 2011); individuals and populations vary genetically and phenotypically within species (Endler 1977), and intra-specific variation can be as great as trait variation across species (Albert et al. 2010; Des Roches et al. 2018). Therefore, models based on the assumption that all members of a species will respond similarly to ecological challenges may be invalid (see Kolbe et al. 2010; Kelly et al. 2012; Valladares et al. 2014; Llewelyn et al. 2016; Moran et al. 2016). In particular, information on the environmental sensitivity of range-edge populations will be vital for understanding future changes in species distributions, since it is at range edges where colonisations and extinctions primarily occur as the climate changes (Valladares et al. 2014; Rehm et al. 2015).

Clinal studies investigating patterns of phenotypic differentiation along environmental gradients can provide insight into how environmental stress has shaped the tolerance of populations (Keller et al. 2013). For example, in anuran amphibians, acid tolerance (Räsänen et al. 2003) and thermal tolerance (Hoppe 1978) vary between populations in a pattern consistent with the cline of each environmental stressor. By quantifying and comparing the sensitivity of populations along environmental clines, estimates of a species’ capacity to adjust, either via phenotypic plasticity or local adaptation (or a combination of both), to altered environmental conditions can be generated (Llewelyn et al. 2016; Pontes-da-Silva et al. 2018). For example, range-edge populations may hold a reservoir of pre-adapted genes that might, with sufficient gene flow, facilitate adaptation at the species level (Aitken and Whitlock 2013). Studies of variation in stress tolerance along environmental clines have demonstrated the capacity for populations to adjust their phenotypes to local conditions (Hoffmann and Harshman 1999; Hoffmann et al. 2002; Arthur et al. 2008; Gilchrist et al. 2008; Sinclair et al. 2012; Keller et al. 2013), and have shown that phenotypic divergence is often adaptive (Rajpurohit and Nedved 2013).

Water availability is a key environmental factor that is particularly important for amphibians. Amphibian skin is highly permeable to water (Young et al. 2005) and thus the ability of anurans to occupy terrestrial habitats is dependent on their ability to resist and avoid desiccation. In adult anurans, hydration state affects signalling behaviours and reproductive success (Mitchell 2001), locomotor performance (Gatten 1987; Hillman 1987), and predator avoidance and the ability to catch prey (Titon et al. 2010). Similarly, water availability has major effects on offspring fitness, particularly in terrestrial-breeding species, as the outer capsule of their eggs is almost completely permeable to water (Bradford and Seymour 1988; Mitchell 2002a; Andrewartha et al. 2008). Terrestrial embryos that develop on relatively dry soils are typically smaller (Mitchell 2002a; Andrewartha et al. 2008; Eads et al. 2012), develop more slowly (Bradford and Seymour 1985), have reduced survival (Martin and Cooper 1972; Bradford and Seymour 1988; Eads et al. 2012) and are more often malformed (Mitchell 2002a; Eads et al. 2012). Thus, low environmental water availability is likely to induce strong directional selection through its negative effects on survival, reproduction and growth. This in turn can potentially lead to genetic or plastic differences in desiccation tolerance among populations that occur across a range of hydric environments. Despite this, there are few studies of population-level variation in anuran desiccation tolerance across clines of water availability (but see Van Berkum et al. 1982), and no studies investigating whether such patterns exist at early developmental stages, where selection on desiccation tolerance should be especially strong given that embryos cannot move to escape unfavourable microclimates.

Here, we quantified intra-specific variation in traits that reflected adult and embryonic desiccation tolerance in populations of the terrestrial-breeding frog *Pseudophryne guentheri*. This species is highly suited to investigating geographic patterns in trait variation, as populations are distributed over a large area of southwestern Australia (Fig. 1) that spans a pronounced rainfall gradient (∼300-1250 mm annual rainfall). Furthermore, *P. guentheri* inhabits a region that has experienced a substantial decline in winter rainfall over the past 40 years (19% reduction since the 1970s) (Smith 2004; IOCI 2012; Andrich and Imberger 2013; CSIRO and Bureau of Meteorology 2016). Rainfall in this region is predicted to decline further in the coming decades (Gallant et al. 2007; Bates et al. 2008; Smith and Power 2014; CSIRO and Bureau of Meteorology 2015) and there is concern that the range of this species will contract (Arnold 1988). We investigated whether adult *P. guentheri* show clinal variation in their water balance and whether the desiccation tolerance of their offspring differs in line with the rainfall gradient. Embryonic stages of terrestrial breeders are particularly vulnerable to desiccation as they cannot escape unfavourable microclimates (Martin and Cooper 1972; Bradford and Seymour 1988; Mitchell 2002a; Eads et al. 2012). Therefore, local precipitation regimes are likely to drive the strength of selection on desiccation tolerance traits in embryos, which may lead to spatial variation in embryonic desiccation tolerance among populations. We focused on quantifying variation in a range of traits putatively tied to fitness, including survival, time to hatch after inundation, wet mass at hatching, hatchling malformations and swimming performance.

**Figure 1.**
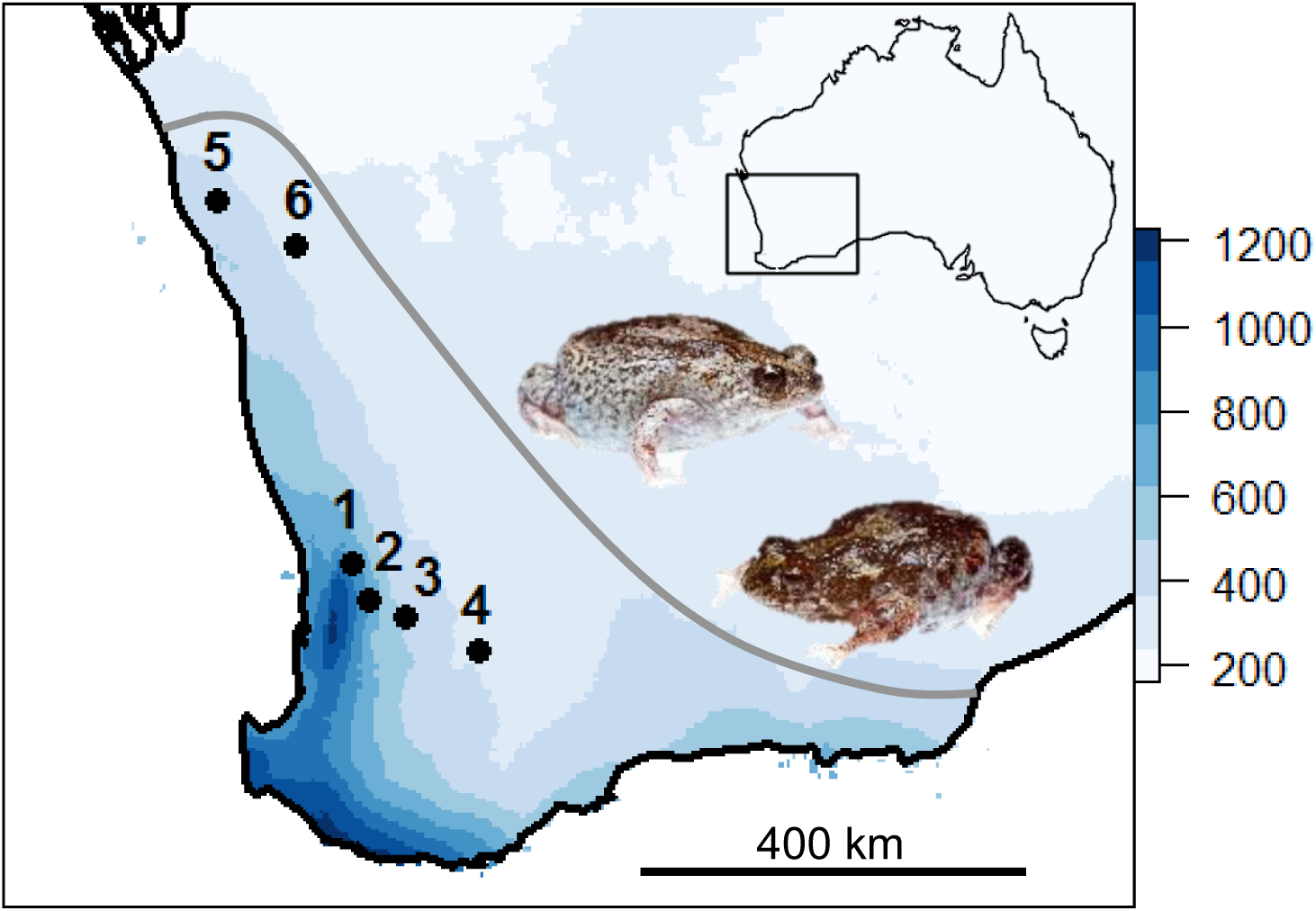
Map showing the distribution of *P. guentheri* in Western Australia (grey line; based on occurrence records from the Atlas of Living Australia) and the location of collection sites, overlaid with annual mean rainfall data (mm). Populations are numbered by increasing aridity. Female *P. guentheri* left, male right.

## MATERIALS AND METHODS

### Ethics statement

All animal experiments were conducted in accordance with the University of Western Australia’s (UWA) Animal Ethics Committee (permit number RA/3/100/1466). Fieldwork was conducted under permit SF010807 issued by the Western Australian Department of Biodiversity, Conservation and Attractions.

### Study species

*Pseudophryne guentheri* (Anura: Myobatrachidae) is a small (26-33 mm snout-to-vent length) terrestrial-breeding frog endemic to the southwest of Western Australia (Tyler and Doughty 2009). Its range abuts that of an inland, arid-adapted member of this widespread genus (*P. occidentalis*). Breeding takes place in autumn and early winter following seasonal rainfall. Males excavate burrows in areas that are likely to be flooded and call from the burrow entrance to attract females (Anstis 2013). After a female has selected and approached a male, mating occurs inside burrows. Females deposit clutches of 60 to 300 eggs directly onto the soil and encapsulated embryos develop terrestrially (Anstis 2013). Hatching occurs when burrows flood in late winter (Eads et al. 2012) and tadpoles complete their development in ephemeral water bodies in about three months (Anstis 2013).

### Animal collection and study sites

Adult *P. guentheri* were collected by hand and in pit-fall traps from six study sites located in the centre and northern limit of the species’ range (Fig 1) in May and June 2016. Sites were distributed across a rainfall gradient, with the wettest site (population 1) receiving ∼790 mm of rain per year and the driest site (population 6) receiving ∼330 mm of rain per year (Table 1). In total, 10 to 19 (mean = 15.8, Table 1) calling males were collected from each site. Gravid females were more difficult to locate and hence our collection of females was restricted to four sites (Table 1). Adult frogs were temporarily housed in small (4.4 L) plastic containers containing moist sphagnum moss and transported to the University of Western Australia in Perth within five days of collection. Frogs were then housed in a controlled-temperature room at 16°C with an 11/13 h light/dark photoperiod to mimic winter conditions, and were fed a diet of pinhead crickets.

**Table 1.** Site characteristics and sample numbers (*N*) and average egg size for each *P. guentheri* population sampled, with populations numbered by increasing aridity.

| Collection site | Population No | Longitude | Latitude | *N* |  |  | Egg diameter<br>± SD (mm) | Number of days between first<br>rainfall* (≥ 5mm/day) and<br>animal collection date (± SD) | Annual mean<br>precipitation<br>(mm) | Annual mean<br>temperature<br>(°C) |
| --- | --- | --- | --- | --- | --- | --- | --- | --- | --- | --- |
|  |  |  |  | ♂ | ♀ | F<sub>1</sub>* |  |  |  |  |
| Chidlow | 1 | 31°53'05.5"S, 116°18'48.0"E |  | 17 | 5 | 647 | 2.576 ± 0.107 | 27 ± 11 | 788 | 17.2 |
| Flint Plot | 2 | 32°17'01.4"S, 116°31'24.1"E |  | 12 | 4 | 453 | 2.389 ± 0.053 | 11 ± 6 | 654 | 16.7 |
| Pingelly | 3 | 32°28'25.9"S, 116°58'27.9"E |  | 19 | 8 | 911 | 2.221 ± 0.150 | 7 ± 5 | 428 | 16.6 |
| Dudinin | 4 | 32°49'17.4"S, 117°53'01.0"E |  | 18 | 0 | 0 | - | 12 ± 4 | 358 | 16.5 |
| Binnu | 5 | 28°02'30.8"S, 114°39'36.0"E |  | 19 | 5 | 870 | 2.165 ± 0.103 | 6 ± 5 | 352 | 19.9 |
| Mullewa | 6 | 28°31'07.3"S, 115°38'11.4"E |  | 10 | 0 | 0 | - | 6 ± 4 | 329 | 20.5 |

\* of the breeding season (May-June 2016)
Note: Climate data were obtained from the Bureau of Meteorology and are interpolated values for the specific coordinates of each population, averaged from 1980 to 2017. \*First generation offspring obtained via within-population parental crosses from four localities.

### Dehydration and rehydration assays in adult males

Rates of dehydration and rehydration were assessed in 90 adult males collected across all six populations, with a minimum sample size of 10 males per population (mean = 15). Frogs were maintained at 16°C for two to five days to reflect winter conditions, during which time they were fed *ad libitum*. Body mass during this period remained stable (± 2.5% body mass). One day prior to commencement of the experiments, all food was removed to minimise weight changes due to defecation during water balance experiments, and sphagnum moss was moistened to ensure that frogs were fully hydrated.

#### Dehydration rate

After the urinary bladder was emptied by cannulation, frogs were weighed (= 100% standard mass) and then placed individually in desiccation cages. Desiccation cages consisted of a PVC ring (diameter = 5.5 cm, height = 2 cm) wrapped in Nylon mesh that allowed frogs to be weighed without handling. Desiccation cages holding frogs were weighed (± 0.01g, GX-600 high precision balance, A&D Weighing, NSW, Australia), and placed into a desiccating chamber containing Drierite (W. A. Hammond, Xenia, OH, USA). Cages were weighed every 30 min until frogs reached 85% of their standard mass. This amount of water loss via evaporation and respiration was reversible and did not cause adverse effects, and frogs moved very little during trials.

#### Rehydration rate

Immediately after the desiccation trials, frogs were removed from the cages and placed in lidded petri dishes containing deionised water to a level of 1 cm. This ensured that a frog’s ventral patch was exposed to the water at all times and minimised their movements. Rehydration occurred rapidly and frogs were blotted dry and weighed every 10 min until regaining 100% of their standard mass.

As rates of water loss and water uptake are dependent on body size (Withers et al. 1982; Wygoda M. 1984; Titon and Gomes 2015), dehydration and rehydration rates are expressed as area specific measurements, calculated for each frog by the equation

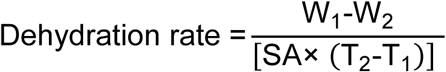

where W1, and W2 indicate initial and final body mass, and T1 and T2 represent start and end time of each trial (Liu and Hou 2012). The same equation was used to calculate area-specific rehydration rates. The surface area of the frogs was estimated as SA (cm^2^) = 9.9 (standard weight)^0.56^ (McClanahan and Baldwin 1969).

### In-vitro fertilisation methods

In-vitro fertilisation was used to perform controlled within-population crosses in the laboratory for four populations where females were obtained (population 1, 2, 3 and 5). The eggs of each female were divided equally into five groups and fertilised separately with sperm from one of five males at random. This allowed us to control for potential parental compatibility effects on offspring fitness, which have been identified in this species (Eads et al. 2012) and in other anurans (Dziminski et al. 2008). Due to the restricted number of sires, we used the same male’s sperm to fertilise several females in some populations (see Statistical Analysis). Males were euthanized via ventral immersion in <0.03% benzocaine solution for 10 min, followed by double pithing. Both testes were removed, blotted dry and weighed to the nearest 0.1 mg (precision balance XS204, Mettler Toledo) and placed on ice. Testes were then macerated in 20 to 615 µL (adjusted according to the weight of the testes) of chilled 1:1 standard amphibian ringer (SAR; 113mM NaCl, 2mM KCl, 1.35 mM CaCl2, and 1.2 mM NaHCO_3_). This buffer has a similar osmolality to the male’s reproductive tract and keeps spermatozoa in an inactive state (Byrne et al. 2015), allowing sperm storage for extended periods of time (days - weeks) without considerable declines in motility (Browne et al. 2001; Kouba et al. 2003). Sperm concentrations in testes macerates were measured using an improved Neubauer haemocytometer (Hirschmann Laborgeräte, Eberstadt, Germany), and sperm suspensions were diluted with 1:1 SAR to produce stock solution of 100 sperm per μl.

Ovulation in females was induced via two subcutaneous injections of the hormone LHRHa over the course of two days (Silla 2011). On day one, a priming dose of 20% of the ovulatory dose was administered, followed by an ovulatory dose (2 μg LHRHa per 1g female standard weight, diluted in 100 μl of SAR) 22 hours after the first injection. Approximately ten hours after administration of the ovulatory dose, eggs were gently squeezed from each female onto a clean surface. They were then moistened with SAR and divided equally among five small petri dishes and placed on ice until fertilisation. Following Eads et al. (2012), a small volume of sperm suspension was pipetted onto one edge of the petri dish, followed by a larger volume of diluted SAR solution (one part SAR to four parts deionised water) which activated the sperm. To promote fertilisation, each dish was manually agitated for 20 sec to mix the solutions and eggs. After 15 minutes, eggs were backlit and photographed (while submerged in water to minimize refraction) using a digital imaging camera (Leica DFC320) attached to a light microscope (Leica MZ7.5) at X 6.3 magnification. These images were used to measure the ovum diameter of a random sample of 50 eggs from each female, using ImageJ software (Abràmoff et al. 2004). One hour after combining eggs and sperm, fertilisation success was scored by counting eggs that had rotated (Gosner Stage 1; Gosner 1960). All fertilisations were initiated between the 12^th^ and 26^th^ of May 2016.

### Soil preparation and embryo incubation

Fertilised eggs were assigned to one of three water-potential (ψ) rearing environments: a wet treatment (ψ = −10 kPa), an intermediate treatment (ψ = −100 kPa) and a dry treatment (ψ = −400 kPa). These water potentials represented a range of hydric conditions in which egg clutches develop in the field, which depends largely on the time since rainfall (N. J. Mitchell, unpubl. data).

Soil water potentials were established by oven-drying homogenised soil, previously collected from a single *P. guentheri* breeding site (Pinjar, see Eads et al. 2012), at 80 °C for 24 hours. Soil was distributed into small, lidded containers (dimensions: 7.5 x 11 x 4 cm, 50 g of soil per container), and re-wetted with an appropriate mass of deionised water using a soil water-retention curve previously determined for the same soil type (N. J. Mitchell, unpubl. data) and allowed to equilibrate. The water content of the soil (g/g of dry soil) was approximately 50% in the wet treatment, 33% in the intermediate treatment, and 21% in the dry treatment.

Eggs were distributed onto soils 7 - 9 hours after fertilisation, with three replicates per water potential treatment and family (sire-by-dam combination). Small plastic rings (nylon plumbing olives, 12 mm in diameter) were placed to surround egg clusters and were labelled to identify individual crosses. The containers housing the eggs and soils were then placed in incubators (model i-500, Steridium, Australia) set at 16 ± 0.5 **°**C and weighed every two days to ensure that soil water potentials remained stable throughout incubation. Embryos were monitored every two days and any mouldy eggs were removed. Embryonic survival was recorded for each family as the percentage of fertilised eggs that hatched (see below).

### Desiccation tolerance of embryos and hatchlings

#### Time to hatching

In the *Pseudophryne genus*, hatching is triggered by the flooding of terrestrial nests following weeks of seasonal winter rainfall. In the absence of flooding, embryos slow their metabolism, halt development and remain dormant until flooding (or death) occurs (Bradford and Seymour 1985; Martin 1999). Thus in this study, hatching was induced by placing embryos individually in small tubes containing 2 ml of deionised water to mimic conditions in the wild. Hatching was induced at 33 days after fertilisation following Eads et al. (2012), who established that emerging *P. guentheri* tadpoles have reached a stage in their development that renders them able to hatch and survive in the rearing habitats (Gosner Stage 26) after 33 days at 16 °C. Embryos were monitored every 30 min, and once a hatchling escaped the egg capsule, the time to hatching was recorded.

#### Swimming performance

Swimming performance was recorded 6 - 12 hours after hatching for a subset of hatchlings (*N* = 556 across all populations, with a minimum sample size of 30 per population and treatment). For this purpose, individual hatchlings were placed in a petri dish (diameter = 15 cm) containing water to a level of 1 cm. After an initial acclimation period of 1 min, a glass cannula was used to gently poke the tail of each hatchling, which elicited a burst swimming response. A video camera (Canon PowerShot G16, recording at 60 fps) installed 30 cm above the petri dish was used to film three burst swimming responses for each hatchling, and their movement was later tracked and analysed using EthoVision v8.5 software (Noldus et al. 2001). EthoVision enabled the quantification of the following swimming parameters: maximum velocity (cm s^-1^), mean velocity (cm s^-1^) and total distance moved (cm). We also recorded mean meander (deg cm^-1^), a measure of the straightness of the swimming response, as dry rearing environments can lead to asymmetrically shaped hatchlings (Eads et al. 2012) that swim in a more circular motion (N. J. Mitchell, pers. observations). A hatchling was considered moving when it exceeded 0.45 cm s^-1^. Since each video recording contained three burst swimming responses with periods of no movement in between them, EthoVision only analysed frames in which a hatchling moved faster than 0.45 cm s^-1^ (consequently merging the three swimming responses for each hatchling). Immediately following the swimming performance trials, hatchlings were euthanized in <0.03% benzocaine and preserved in 10% neutral buffered formalin.

#### Hatchling wet weight and malformations

Wet masses of preserved hatchlings were recorded to the nearest 0.001 g after blotting on tissue. Hatchlings were then photographed in lateral view while submerged in water using a digital imaging camera (Leica DFC320) attached to a light microscope (Leica MZ7.5) at X 6.3 magnification. These images were used to score any malformations for each hatchling. Malformations consisted of three forms of notochord malformations (scoliosis – lateral curvature of the spine; lordosis – concave curvature of the spine; and kyphosis – convex curvature of the spine) and edemas (abnormal accumulation of fluids in tissues) (Fig. 2). Collectively, these deformities represented 84% of the total number of malformations observed. Malformations were quantified for each combination of treatment and sire-by-dam family as the percentage of hatchlings that showed a malformation of any kind.

**Figure 2.**
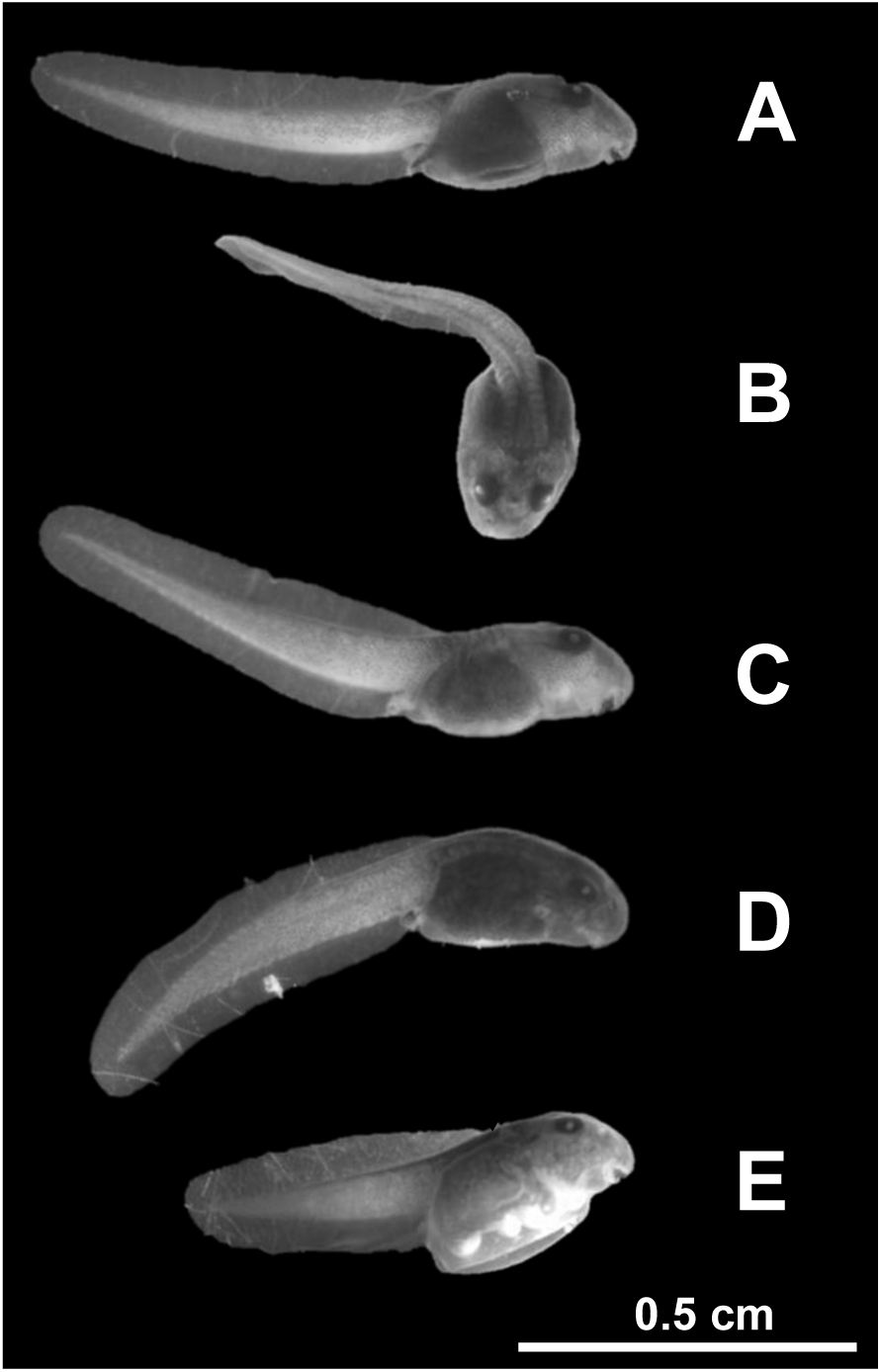
Types of malformations in *P. guentheri* hatchlings scored in this study: (A) normal shaped hatchling without malformations, (B) hatchling with scoliosis, (C) hatchling with lordosis, (D) hatchling with kyphosis and (E) hatchling with edema.

### Statistical analysis

All analyses were performed in R version 3.4.3 (R Development Core Team 2017). To ensure that data complied with assumptions of normality, Q-Q plots of all residuals were inspected prior to analysis.

#### Adult de- and rehydration rates

Rehydration rate was log_10_-transformed and the rate of dehydration was transformed using the Box-Cox method (Box and Cox 1964) to fulfil assumptions of normality. We then used one-way ANOVAs, followed by Tukey HSD post-hoc tests, to test whether dehydration and rehydration rates significantly differed between populations. Linear regression analyses were used to test for relationships between dehydration or rehydration rates and mean annual rainfall.

#### Desiccation tolerance of embryos and hatchlings

For four populations where we created within-population crosses we used linear mixed-effects models (with restricted maximum-likelihood methods; REML) to investigate variation in offspring fitness traits resulting from the water potential treatment. Hatchling wet weight, time to hatching, total distance moved, and mean meander were transformed using the Box-Cox method (Box and Cox 1964) to fulfill assumptions of normality. Linear mixed-effect models were run using the lme4 package in R (Bates et al. 2015), with treatment, population and the population-by-treatment interaction treated as fixed effects and dam fitted as a random effect. We also added sire identity as a random effect to all models to control for the repeated use of sperm donors across IVF trials. Ovum size was added as a covariate in all analyses to control for some trait variation due to maternal effects (Eads et al. 2012). The significance levels of fixed effects were evaluated using Wald chi-squared tests on the full model.

Fertilization rates, survival and malformation data were binomial variables and thus a generalized linear mixed-effects model (GLMM) with a logit-link function was used for the analysis of these traits. Fixed effects were treatment, population and the population-by-treatment interaction, and sire and dam were added as random effects. The significance of the fixed effects of treatment and population and their interaction, were evaluated using Wald Z tests, and 95% Wald confidence intervals were obtained using the “confint” function from the MASS package in R (Venables and Ripley 2002).

## RESULTS

### Dehydration and rehydration rates

Across all six study populations, males lost 15% of their standard mass on average after 4.98 ± 1.39 SD h (range: 2.03 - 8.20 h) in the desiccation chamber, and on average rehydrated from this benchmark to full hydration in less than an hour (0.82 ± 0.39 SD h, range: 16 - 120 min). Time to dehydrate and rehydrate correlated strongly with body size (*P* < 0.001), and therefore only area-specific dehydration and rehydration rates are used to compare populations (see Materials and Methods). Rates of dehydration differed significantly among populations (F_5,84_=11.58, *P* < 0.001), with the highest in males from the wettest site (population 1) and a trend towards decreasing rates with increasing aridity (Fig. 3A). Rehydration rates followed a similar pattern (Fig 3B), but the population effect was not significant (F_5,84_ = 0.964, *P* = 0.445).

**Figure 3.**
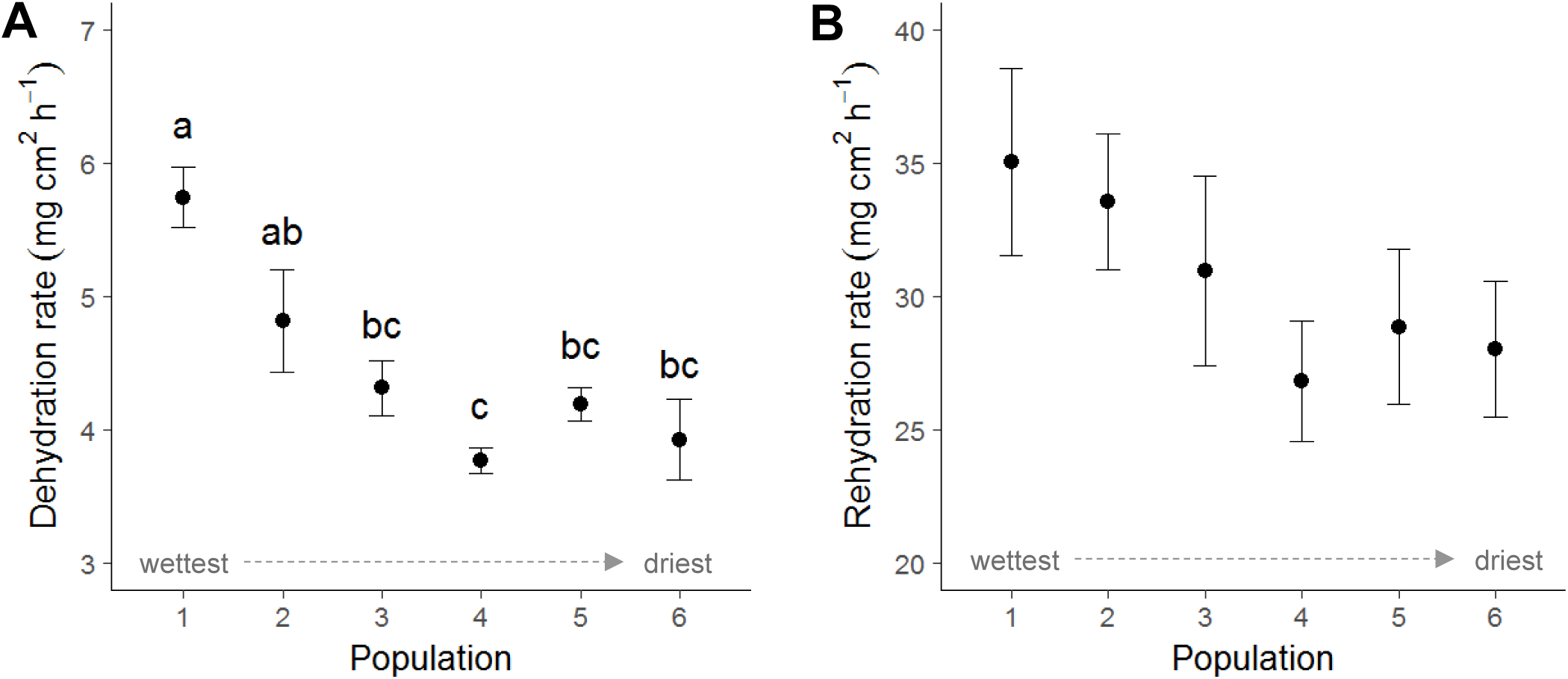
Variation in (A) dehydration and (B) rehydration rates (mean ± SE) among the six *P. guentheri* populations located along a natural rainfall gradient (790 - 330 mm/year). The x-axis lists populations from wettest to driest sites. Populations that do not share the same letters are significantly different (Tukey HSD post-hoc tests, *P* < 0.05).

### Desiccation tolerance of embryos and hatchlings

#### Survival

Embryonic survival on wet soils (high water potential) was high (> 93%) in all four populations (Fig. 4A). Lower soil water potentials negatively affected survival rates, although the severity of the treatment effect varied among populations, as evident from significant population-by-treatment effects (Table 2). The population from the wettest site (population 1) showed the largest reduction in survival in response to dry and intermediate soil moisture, and this difference tended to decrease in populations originating from sites of increasing aridity (Fig. 4A). As such, low soil water potentials had very little effect on embryonic survival in the driest population (population 5), with over 88% of embryos surviving to hatching stage.

**Figure 4.**
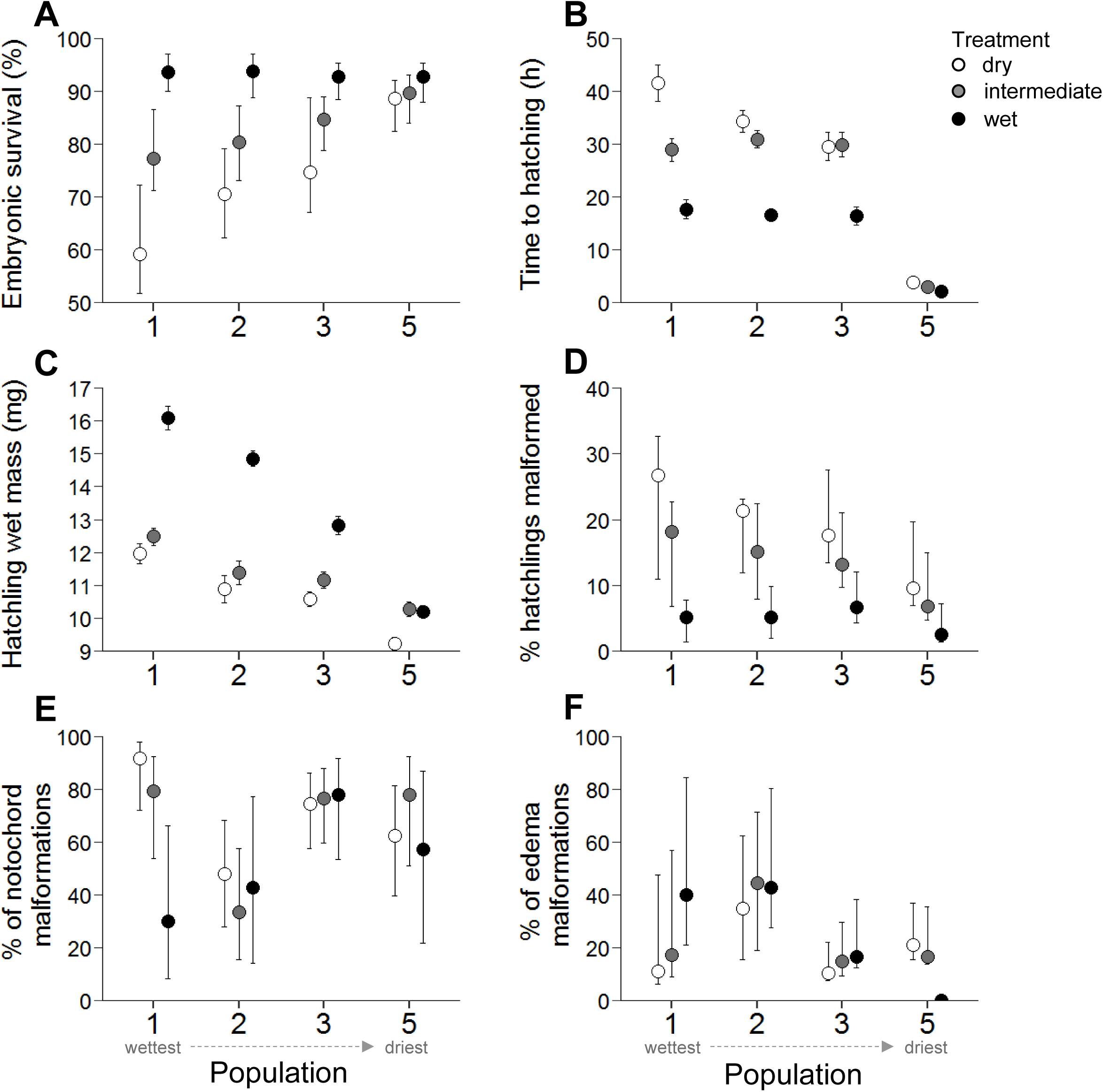
Embryonic and larval *Pseudophryne guentheri* trait responses (mean with 95% confidence intervals) for four populations, reared on soil at three water potentials; dry (white circles), intermediate (grey circles) and wet (black circles). The x-axis shows populations arranged from the wettest (1) to the driest (5) sites. (A) embryonic survival (proportion of fertilised eggs that hatched), (B) time to hatching (h), (C) wet weight at hatching (mg), (D) proportion of malformed hatchlings, (E) proportion of malformations in the notochord category and (F) proportion of malformations in the edema category.

**Table 2.** Mixed-effects model results of embryonic and larval *P. guentheri* traits associated with desiccation tolerance

| **Trait** | **N** | **Source** | **df** | **X<sup>2</sup>** | **P** | **Sig.** |
| --- | --- | --- | --- | --- | --- | --- |
| Embryonic survival (%) | 2844 | Treatment | 2 | 98.398 | < 0.001 | *** |
|  |  | Population | 3 | 8.874 | 0.031 | * |
|  |  | Population x Treatment | 6 | 19.924 | 0.003 | ** |
|  |  | Ovum size | 1 | 0.926 | 0.336 | ns |
| Time to hatching (h) | 2378 | Treatment | 2 | 745.596 | < 0.001 | *** |
|  |  | Population | 3 | 104.731 | < 0.001 | *** |
|  |  | Population x Treatment | 6 | 173.249 | < 0.001 | *** |
|  |  | Ovum size | 1 | 0.455 | 0.499 | ns |
| Wet weight at hatching (mg) | 2344 | Treatment | 2 | 864.308 | < 0.001 | *** |
|  |  | Population | 3 | 5.641 | 0.130 | ns |
|  |  | Population x Treatment | 6 | 295.199 | < 0.001 | *** |
|  |  | Ovum size | 1 | 3.770 | 0.05 | * |
| Proportion of hatchlings malformed | 2382 | Treatment | 2 | 59.152 | < 0.001 | *** |
|  |  | Population | 3 | 5.934 | 0.115 | ns |
|  |  | Population x Treatment | 6 | 2.940 | 0.816 | ns |
|  |  | Ovum size | 1 | 2.545 | 0.111 | ns |
| Proportion of notochord malformations | 2382 | Treatment | 2 | 2.929 | 0.231 | ns |
|  |  | Population | 3 | 18.234 | < 0.001 | *** |
|  |  | Population x Treatment | 6 | 12.272 | 0.056 | ns |
|  |  | Ovum size | 1 | 0.006 | 0.939 | ns |
| Proportion of edema malformations | 2382 | Treatment | 2 | 2.811 | 0.245 | ns |
|  |  | Population | 3 | 13.673 | 0.003 | ** |
|  |  | Population x Treatment | 6 | 2.350 | 0.885 | ns |
|  |  | Ovum size | 1 | 1.489 | 0.222 | ns |

#### Time to hatching

Across all populations, the average time that the terrestrial embryos required to hatch after submergence in water significantly increased with decreasing soil water potentials (Table 2, Fig. 4B). That is, embryos reared on relatively wet soils hatched more quickly than their siblings reared on drier soils. However, there were marked differences in hatching times between populations, and the way treatment affected time to hatching also differed significantly among populations (Table 2, Fig. 4B). As such, hatchlings originating from the driest population (population 5) hatched significantly more quickly than hatchlings from the other three populations, irrespective of the rearing environment (Fig. 4B). Hatchlings reared on wet soils from populations 1, 2 and 3 hatched after approximately 17 hours, but required approximately 30 hours when reared at intermediate water potentials. Furthermore, hatchlings originating from the wettest site (population 1) took > 40 h to hatch, but this increase in hatching time was less substantial in population 2, and populations 3 and 5 exhibited similar hatching times in dry and intermediate rearing environments (Fig. 4B).

#### Wet mass at hatching

Low soil water potentials significantly reduced the wet mass of hatchlings. Similar to the pattern observed in the previous two traits, the population-by-treatment interaction was significant, suggesting that the severity of the treatment effect varied among populations (Table 2). Hatchlings from the wettest site (population 1) showed the largest reduction in hatchling size in response to low soil water potentials, and this decline in size decreased in populations originating from sites with increasing aridity (Fig 4C). Accordingly, hatchling wet mass from the driest site (population 5) hardly varied between dry, intermediate or wet rearing environments (Fig. 4C). As expected, ovum size significantly affected the wet mass of hatchlings (Table 2).

#### Malformations

The total percentage of malformed hatchlings was low (∼ 5%) in the wet treatment in all four populations (Fig 4D) but increased with decreasing soil water potential. Malformation rates did not differ significantly between populations, and there were no significant population-by-treatment interactions for this trait. Soil water potential treatments did not significantly affect the relative occurrence of notchord or edema type malformations, but there were significant population effects for both these traits (Table 2, Fig. 4D).

#### Swimming performance

Low soil water potential significantly reduced the swimming performance of hatchlings (Table 3), with maximum and mean swimming velocity decreasing (Fig. 5A & B) and the total distance moved declining in dry and intermediate rearing environments (Fig. 5D, Table 3). Furthermore, mean meander increased in response to low soil water potentials (Fig. 5C), indicating that hatchlings swam more linearly when reared in wetter conditions. The effect of the soil water potential treatment on swimming performance differed significantly among populations (Table 3), with the wettest population (population 1) showing the largest declines in velocity and largest increases in mean meander in response to the dry and intermediate treatment. In contrast, swimming performance in hatchlings from the driest population (population 5) was hardly affected by the dry treatment (Fig. 5)

**Figure 5.**
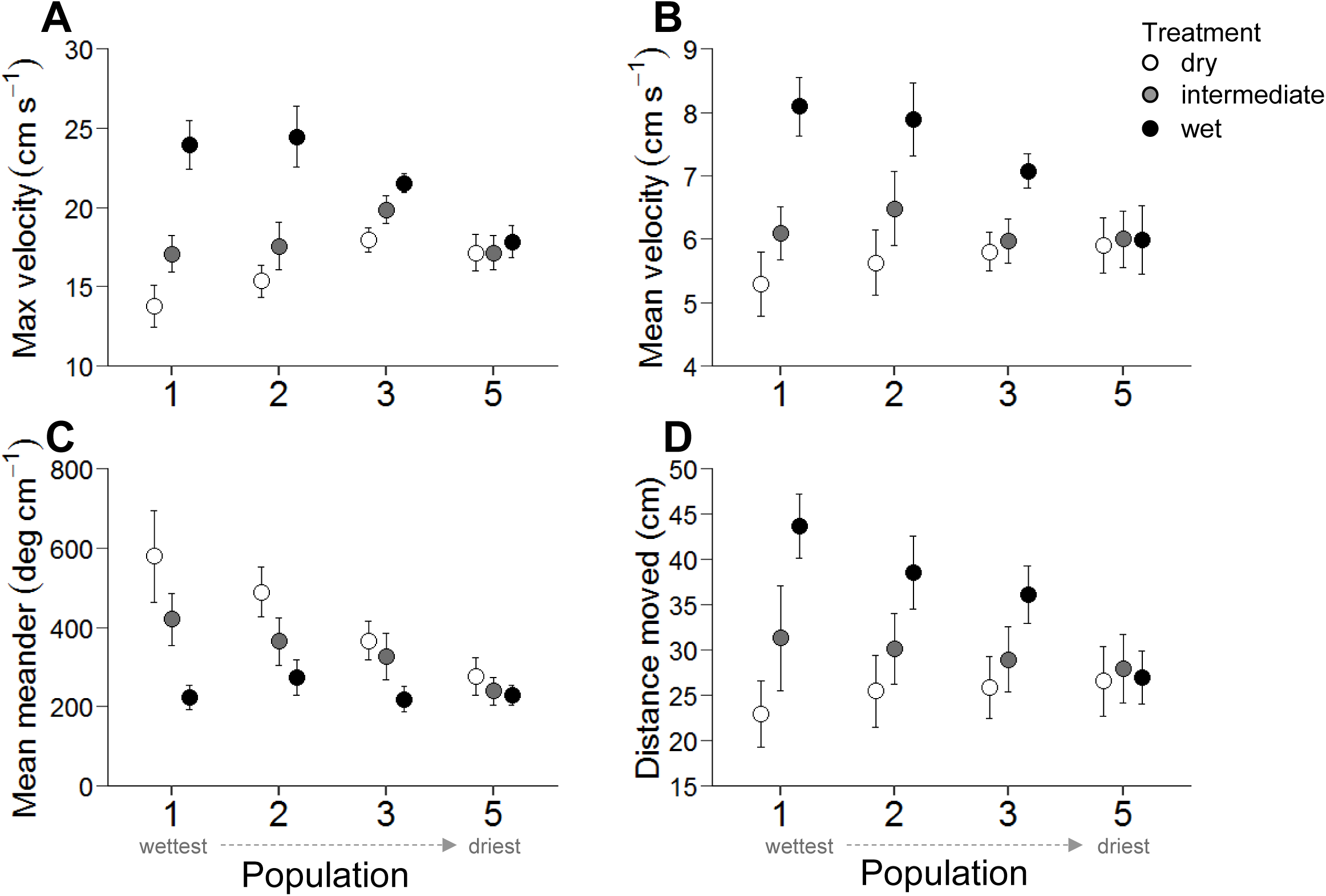
Swimming performance (mean with 95% confidence intervals) of *P. guentheri* hatchlings from four populations, reared on soil at three water potentials; dry (white circles), intermediate (grey circles) and wet (black circles). The x-axis shows populations arranged from the wettest (1) to the driest (5) sites. (A) maximum velocity (cm s^-1^), (B) mean velocity (cm s^-1^), (C) mean meander (deg cm ^-1^) and (D) total distance moved (cm).

**Table 3.** Mixed-effects model results of swimming performance traits in *P. guentheri* hatchlings originating from four populations along a natural rainfall gradient.

| Trait | N | Source | df | X<sup>2</sup> | P | Sig. |
| --- | --- | --- | --- | --- | --- | --- |
| Maximum velocity (cm s<sup>-1</sup>) | 556 | Treatment | 2 | 197.270 | < 0.001 | *** |
|  |  | Population | 3 | 12.499 | 0.006 | ** |
|  |  | Population x Treatment | 6 | 95.770 | < 0.001 | *** |
|  |  | Ovum size | 1 | 0.4347 | 0.510 | ns |
| Mean velocity (cm s<sup>-1</sup>) | 556 | Treatment | 2 | 197.270 | < 0.001 | *** |
|  |  | Population | 3 | 12.499 | 0.006 | ** |
|  |  | Population x Treatment | 6 | 95.770 | < 0.001 | *** |
|  |  | Ovum size | 1 | 0.4347 | 0.510 | ns |
| Mean meander (deg cm<sup>-1</sup>) | 556 | Treatment | 2 | 138.066 | < 0.001 | *** |
|  |  | Population | 3 | 11.747 | 0.008 | ** |
|  |  | Population x Treatment | 6 | 25.783 | < 0.001 | *** |
|  |  | Ovum size | 1 | 4.518 | 0.036 | * |
| Total distance moved (cm) | 556 | Treatment | 2 | 66.881 | < 0.001 | *** |
|  |  | Population | 3 | 5.699 | 0.127 | ns |
|  |  | Population x Treatment | 6 | 17.385 | 0.008 | ** |
|  |  | Ovum size | 1 | 1.765 | 0.184 | ns |

## DISCUSSION

Our findings demonstrate significant intra-specific variation in traits related to desiccation tolerance in *Pseudophryne guentheri* (adult males and first generation offspring), consistent with patterns of genetic adaptation and/or phenotypic plasticity in response to local water availability. These results emphasize the importance of considering population variation in fitness-related traits when making predictions about the fate of species in the face of climate change. We discuss these key findings in turn below.

### Dehydration and rehydration rates in adult males

Our observation that populations significantly differed in their water balance (dehydration and rehydration rates) was in accordance with annual variation in rainfall in these populations (Fig. 3). Dehydration negatively affects survival and fitness in anurans through its effects on locomotor performance (Claussen 1974; Gatten 1987; Moore and Gatten 1989; Köhler et al. 2011), which can impede predator escape, foraging behaviour, territorial defence, signalling and mating (Mitchell 2001; Titon and Gomes 2015). Consequently, lower rates of dehydration are likely to benefit frogs occupying dry habitats, as they allow individuals to leave moist retreats for longer to perform ecologically important behaviours (Feder and Londos 1984; Winters and Gifford 2013). While our experiments were not designed to determine whether variation in dehydration rates was a consequence of genetic adaptation or phenotypic plasticity, our results reflect interspecific patterns for amphibians reported elsewhere. For example, in anurans, resistance to evaporative water loss (EWL) correlates with environmental water availability (Wygoda M. 1984; De Andrade and Abe 1997), with terrestrial specialists exhibiting lower rates of EWL than species occupying primarily aquatic habitats (Young et al. 2005). Similarly, Winters and Gifford (2013) found variation in EWL rates at the population level, with populations of lungless salamander *(Plethodon montanus*) from dry, low-elevation areas dehydrating more slowly relative to those from wetter, higher-elevation areas.

One finding less consistent with the current understanding of amphibian water balance was our observation that mean annual rainfall and rates of rehydration had a significant positive relationship. That is, males from xeric sites rehydrated more slowly than males from mesic sites. This finding is in contrast to interspecific patterns, where rehydration rates are generally highest in species from arid environments (Ewer 1952; Bentley et al. 1958; Dumas 1966; Van Berkum et al. 1982; Tingley and Shine 2011; Titon and Gomes 2015). Similarly, the single study we found that investigated rehydration rates of an anuran species at the population level demonstrated higher rates of water uptake in populations from drier habitats (Van Berkum et al. 1982). The ability to absorb water rapidly can extend the time that a frog can be active by decreasing the amount of time spent in a refuge for rehydration (Van Berkum et al. 1982), suggesting that fast and efficient hydration is likely beneficial for frogs, particularly those inhabiting dry areas.

One explanation for the slower rehydration rates observed in the more xeric populations could be that the mechanisms that reduce evaporative water loss also reduce water uptake. The thickness of the epidermis (stratum corneum), for example, affects both dehydration and rehydration rates in anurans (Toledo and Jared 1993). There is also evidence that skin thickness is associated with habitat aridity at the species level (Toledo and Jared 1993), with species from drier areas producing a thicker epidermis which reduces dehydration rates whilst also impeding water uptake. Further, aquaporins (water channel proteins) embedded within the pelvic patch (Kubota et al. 2006) can influence rehydration rates in anurans, and variation in the density and type of aquaporins among species has been interpreted as an adaptation of frogs to their respective hydric environment (Suzuki et al. 2007; Ogushi et al. 2010). It is currently unknown whether similar patterns of morphological or physiological differences in the skin properties exist at the population level, and we encourage future studies to explore the underlying mechanisms that drive intra-specific differences in dehydration and rehydration rates across hydric gradients.

An alternative explanation for our rehydration rate data is that our method of exposing the ventral patch of dehydrated frogs to water may not have adequately reflected natural behaviours. Terrestrial frogs rarely rehydrate in free standing water, and instead water uptake is most often achieved via exposure of the ventral patch to moist soil (Wells 2007). Whilst our design minimised potential behavioural co-factors that could influence rehydration rates (e.g. frogs rehydrating on soil could employ a range of tactics, including burying and/or adopting different postures that aid water uptake), future work comparing rates of water uptake in water and moist soil would be beneficial for an evaluation of the generality of the rehydration patterns obtained here.

### Desiccation tolerance of embryos and hatchlings

We found substantial intra-specific variation in traits related to desiccation tolerance in *P. guentheri* embryos and hatchlings. Low soil water potentials generally reduced the expression of traits putatively linked to fitness, although the severity of this effect varied greatly among populations, with populations from the wettest site (pop. 1) being the most negatively affected by dry rearing environments. Embryonic survival, for example, was reduced by ∼35% in the dry treatment in the wettest population, but only by ∼5% in the population originating from the driest site (Fig. 4A). Whilst negative effects of desiccation on the survival of terrestrial embryos are well established (Martin and Cooper 1972; Bradford and Seymour 1988; Mitchell 2002a; Eads et al. 2012), the present study is, to our knowledge, the first to report such marked differences among populations. Nevertheless, we are only able to speculate about the mechanisms driving such intra-specific differences in embryonic survival across the dry rearing environments. One possible explanation is that the structural composition of the egg jelly coat may differ among populations. For example, females from drier areas may produce eggs with thicker capsules, which would limit water loss during embryonic development (Martin and Cooper 1972). Alternatively, there may be physiological processes that enhance desiccation tolerance of embryos developing on drier soils. For example, embryos may slow their metabolism and development rates and extend the period of subsistence on energy reserves to reduce their water requirements (Bradford and Seymour 1985; Podrabsky et al. 2010). We eagerly anticipate future studies designed to uncover the mechanisms underlying variation in desiccation resistance among populations.

We found marked intra-specific differences in the time an embryo was able to hatch after submerging it in water. In particular, embryos from the driest population (pop. 5) hatched quickly, irrespective of the water potential of the soil on which they were reared, whereas time to hatching was prolonged in populations originating from areas with increasing annual precipitation (Fig 4B). Faster hatching times are likely beneficial, as they can allow hatchlings to be washed into stable pools of standing water, where they can feed and seek refuge (Warkentin 2011*a*). This response may be particularly vital in areas where annual precipitation is low and where ephemeral water bodies also dissipate more quickly due to higher ambient temperatures. At the same time, immediate hatching after submergence is not always advantageous. For example, embryos may delay hatching if they are too premature to be fully competent as free-swimming tadpoles (Warkentin 2011*a*).

The strong intra-specific variation in hatching time raises questions about the mechanisms driving these differences. Hatching in amphibians is highly plastic (Warkentin 2011*a*) and can be achieved through chemical or mechanical processes, or a combination of both (Duellmann and Trueb 1986). The chemical hatching mechanism is more common (Warkentin 2011*a*), and embryos of many species have hatching glands which release proteolytic enzymes in response to hatching cues (such as predators or low PO_2_) that digest components of the egg (Carroll and Hedrick 1974; Nokhbatolfoghahai and Downie 2007). It is possible that embryos produced from drier sites had denser hatching glands, synthesised a larger amount of enzyme or released enzymes from vesicles more rapidly (Nokhbatolfoghahai and Downie 2007; Kawaguchi et al. 2009; Cohen et al. 2016). Alternatively, intra-specific differences in hatching time could potentially arise due to variation in development rate. Development rate is negatively affected by low soil water potentials (Bradford and Seymour 1985; Packard 1991) and it is possible that embryos from mesic populations were too premature to hatch after 33 days when reared on dry soils. For example, hatching glands may not have developed, or embryos may have lacked the sensory capacities to perceive the hatching cue (Warkentin 2011*b*). However, hatching glands in Myobatrachids develop relatively early (from Gosner stage 17) and are fully present in embryos by Gosner stage 19 (Anstis 2010), making it unlikely that any slight differences in the maturity of embryos in our study affected their ability to hatch.

A further potential driver of intra-specific differences in hatching time may be the structural composition of the egg capsule. For example, egg capsules from populations occupying drier habitats may swell more rapidly when they rehydrate when flooded, resulting in mechanical rupture of the perivitelline membrane, or an outer egg capsule layer, and so releasing the embryo (Duellmann and Trueb 1986). While understanding the mechanisms driving differences in hatching time was not the focus of our study, in light of our findings we suggest that *P. guentheri* offers an ideal experimental system to explore this further.

Low soil water potentials reduced the wet mass of hatchlings and increased the occurrence of hatchling malformations, in line with earlier studies (Taigen et al. 1984; Mitchell 2002a; Andrewartha et al. 2008; Eads et al. 2012). Embryos were preserved 6 - 12 hours after hatching, and therefore differences in wet weight are unlikely to be the result of variation in body water content as hatchlings had ample time to hydrate. Retarded growth on drier soils may be a consequence of reduced metabolic rates (Bradford and Seymour 1985; Mitchell 2002a) and yolk consumption (Packard 1999). However, wet mass at hatching was only reduced slightly in hatchlings from the driest site when reared on dry soils, whereas wet mass decreased significantly in hatchlings originating from more mesic sites (Fig. 4C). Malformations are associated with insufficient swelling of the egg capsule on dry substrates, which leads to embryos being unable to rotate freely within the perivitelline space and sometimes adhering to the perivitelline membrane (Bradford and Seymour 1988; Mitchell 2002a; Andrewartha et al. 2008). Tadpole malformations can persist after metamorphosis (Plowman et al. 1994) and are therefore likely to have long-term consequences for survival and fitness.

The effect that water potential treatment had on body size and shape explained differences in swimming performance among hatchings from different populations, with hatchlings swimming more slowly, less straight and moving a smaller distance if they were reared on drier soils (Fig 5). Decreased swimming performance is likely to have negative fitness consequences, as rapid swimming allows tadpoles to escape predators and generally aids foraging (Webb 1986; Watkins 1996; Wilson and Franklin 1999; Teplitsky et al. 2005; Walker et al. 2005; Langerhans 2009). As observed for other traits studied here, there was pronounced intra-specific variation in the effects of dry soils on hatchling swimming performance. Swimming performance was unaffected by the dry or intermediate soil water potential treatments in the driest population (pop. 5), but decreased steeply in more mesic populations in response to incubation on dry soils. Interestingly, hatchlings from the driest population swam comparatively poorly, even though other effects of the dry rearing environments were minimal. One explanation could be that egg capsules from mesic populations are more permeable to water, and consequently swell relatively more on moist soils while also dehydrating more rapidly on dry soils. Thus, the benefit of moist soils may be greater in mesic populations. Alternatively, there may be selection for smaller body size in hatchlings from drier sites (Fig. 4C), enhancing their survival via a reduction of deleterious malformations due to lesions with the perivitelline membrane, but at a cost to swimming performance.

One aspect not addressed in this study are behaviours that breeding pairs could employ to aid successful incubation of their offspring within their respective environments. For example, *P. guentheri* males that breed in drier soils might dig deeper burrows which would provide developing embryos with lower (wetter) and more stable soil water potentials. Concurrently, females might preferentially choose to mate with males that are located within high quality oviposition sites, as has been shown in the related species *Pseudophryne bibroni*, where males occupying wetter nest sites had higher mating success (Mitchell 2001). In terrestrial-breeders, other nest traits preferred by females include chamber depth and complexity in the ornate nursery frog (*Cophixalus ornatus*), which better protect egg clutches from biotic or abiotic disturbance (Felton et al. 2006), and nest breadth and vegetation type, which enhances clutch oxygenation in the moss frog *Crinia nimbus* (Mitchell 2002b, 2003). As behavioural selection of microhabitats can play a fundamental role in buffering the strength and direction of selection in natural populations (Doody et al. 2006; Woods et al. 2015; Mitchell and Bergmann 2016; Li et al. 2018), we encourage investigation of geographic patterns of behavioural variation in amphibians, particularly with regards to nest site selection.

In summary, we show significant intra-specific variation in traits associated with desiccation tolerance in *P. guentheri* adults, and in the early developmental stages of their offspring. While dry rearing environments generally had a negative effect on traits putatively linked to fitness, the severity of treatment effects varied greatly among populations in a pattern consistent with the cline in annual precipitation. Together our findings suggest that water availability has influenced patterns of population variation in desiccation tolerance in *P. guentheri*, mediated either by phenotypic plasticity, local adaptation, or a combination of both. Whether the species will decline in response to declines in annual rainfall in southwestern Australia remains unclear. Prior work (Eads et al. 2012) revealed that at least one *P. guentheri* population lacks sufficient additive genetic variation in desiccation tolerance to adapt to changes in water availability. Depending on the generality of this result, the species may have limited capacity to adapt to new abiotic environments (e.g. Kellermann et al. 2009). Yet, if the intra-specific variation in desiccation tolerance we observed in this study is a result of local adaptation, then dry-adapted populations likely possess a reservoir of genes that may facilitate adaptive responses to changes in water availability (Cummins et al. 2019). As many *P. guentheri* populations are now isolated due to extensive habitat fragmentation (Arnold 1988; Hobbs 1993, Cummins et al. 2019), limiting gene flow, the dry-adapted populations could be used as source populations for assisted, targeted gene flow (Aitken and Whitlock 2013) in future efforts to conserve mesic populations. With climate change progressing rapidly, there is increasing focus on conservation of range-edge populations that may harbour the bulk of a species’ adaptive variation (e.g. Rehm et al. 2015; Macdonald et al. 2017). Consequently, the utilisation (e.g. through assisted gene flow) of adaptive variation present in peripheral populations may be vital for reducing extinctions linked to climate change.

## ACKNOWLEDGEMENTS

This work would not have been possible without great support from Stewart Macdonald who located many populations and collected breeding adults, and Brighton Downing who provided extensive assistance in the field and laboratory. We further thank our volunteers who assisted with collecting adults from breeding choruses: Marcus Lee, J. P. Lawrence, Callum Donohue, Blair Bentley and Savannah Victor. This research was supported by funding from the School of Biological Sciences at the University of Western Australia, the ANZ Holsworth Wildlife Research Endowment and the Australian Government’s National Environmental Science Programme through the Threatened Species Recovery Hub. T. Rudin-Bitterli was supported by the International Postgraduate Research Scholarship and the C.F.H. & E.A. Jenkins Postgraduate Research Scholarship. This preprint has been reviewed and recommended by Peer Community In Evolutionary Biology (https://doi.org/10.24072/pci.evolbiol.100079).

